# Transcription factor EB (TFEB) interaction with RagC is disrupted during Enterovirus D68 Infection

**DOI:** 10.1101/2024.03.23.586376

**Authors:** Alagie Jassey, Noah Pollack, Michael A. Wagner, Jiapeng Wu, Ashley Benton, William T. Jackson

## Abstract

Enterovirus D68 (EV-D68) is a picornavirus associated with severe respiratory illness and a paralytic disease called acute flaccid myelitis in infants. Currently, no protective vaccines or antivirals are available to combat this virus. Like other enteroviruses, EV-D68 uses components of the cellular autophagy pathway to rewire membranes for its replication. Here, we show that transcription factor EB (TFEB), the master transcriptional regulator of autophagy and lysosomal biogenesis, is essential for EV-D68 infection. Knockdown of TFEB attenuated EV-D68 genomic RNA replication but did not impact viral binding or entry into host cells. The 3C protease of EV-D68 cleaves TFEB at the N-terminus immediately post-peak viral RNA replication, disrupting TFEB-RagC interaction and restricting TFEB transport to the surface of the lysosome. Despite this, TFEB remained mostly cytosolic during EV-D68 infection. Overexpression of a TFEB mutant construct lacking the RagC binding domain, but not the wild-type construct, blocks autophagy and increases EV-D68 nonlytic release in H1HeLa cells, but not in autophagy defective ATG7 KO H1HeLa cells. Our results identify TFEB as a vital host factor regulating multiple stages of the EV-D68 lifecycle and suggest that TFEB could be a promising target for antiviral development against EV-D68.

**Importance:** Enteroviruses are among the most significant causes of human disease. Some enteroviruses are responsible for severe paralytic diseases such as poliomyelitis or acute flaccid myelitis (AFM). The latter disease is associated with multiple non-polio enterovirus species, including Enterovirus D68 (EV-D68), Enterovirus 71 (EV-71), and coxsackievirus B3 (CVB3). Here we demonstrate that EV-D68 interacts with a host transcription factor, TFEB, to promote vRNA replication and regulate egress of virions from cells. TFEB was previously implicated in viral egress of CVB3, and the viral protease 3C cleaves TFEB during infection. Here we show EV-D68 3C protease also cleaves TFEB after the peak of vRNA replication. This cleavage disrupts TFEB interaction with the host protein RagC, which changes the localization and regulation of TFEB. TFEB lacking a RagC-binding domain inhibits autophagic flux and promotes virus egress. These mechanistic insights highlight how common host factors affect closely related, medically-important viruses differently.

## Introduction

Enteroviruses are ubiquitous pathogens that infect humans and animals, significantly threatening global public health. Enterovirus D68 (EV-D68), an enterovirus that causes acute respiratory illness in infants, was discovered in 1962 but was rarely detected (1–3). However, since 2014, the virus has become increasingly associated with both severe respiratory disease and a neurological condition known as acute flaccid myelitis, which causes limb paralysis in children. No preventative vaccines or specific antiviral treatments are available to treat EV-D68 infection.

EV-D68 belongs to the *Piconarviridae* family of positive sense single-strand RNA viruses, with a genome size of about 7500 nucleotides, encoding a single polyprotein post-translationally processed to structural and nonstructural proteins (4). Like other enteroviruses, infection with EV-D68 leads to an extensive reconfiguration of cellular membranes, and double-membrane autophagosome-like vesicles, which support various phases of the viral life cycle, including genomic RNA replication and nonlytic release, can be observed late during EV-D68 infection (5, 6).

Autophagy is a conserved cellular catabolic process in metazoa that maintains cellular homeostasis by enveloping dysfunctional cytoplasmic contents, including damaged/spent organelles, and targeting them to the lysosomes for degradation and recycling (7). As a basal housekeeping process, autophagy is constitutive at a relatively low level but upregulated in response to various types of stress, such as hypoxia, amino acid starvation, and viral infections (8, 9).

Autophagy has traditionally been thought to be primarily regulated by cytosolic processes, partly due to the observation that enucleated cells can form autophagosomes. However, accumulating evidence now points to an essential role of transcription in regulating stress-induced autophagy, as epitomized by the identification of Transcription Factor EB (TFEB) as the master transcriptional regulator of autophagy and lysosomal biogenesis (10–12). TFEB is a member of the microphthalmia family of basic helix–loop–helix–leucine–zipper (bHLH-Zip) transcription factor (12–15). The subcellular localization of TFEB is regulated by phosphorylation and dephosphorylation. Under basal growth conditions, the Rag GTPases physically interact with TFEB, recruiting it to the surface of the lysosomes, where TFEB is phosphorylated and retained in the cytosol. Upon cellular stress, such as amino starvation or lysosomal damage, TFEB is dephosphorylated and translocates to the nucleus to initiate the transcription of autophagy and lysosomal biogenesis genes (16).

Many enteroviruses are known to trigger autophagic signaling and use components of the autophagic pathway for their benefit. However, only CVB3 has been shown to utilize TFEB for its egress (17). TFEB is cleaved by the 3C protease of CVB3, and the cleaved form has its own activity in promoting viral egress (17). Whether other enteroviruses similarly utilize TFEB is unclear. Here, we demonstrate that TFEB is essential for EV-D68 infection. Knockdown of TFEB reduces EV-D68 genomic RNA replication without significantly impacting viral binding or entry into host cells. We show that the EV-D68 3C protease cleaves TFEB at the N-terminus immediately after peak viral RNA replication, much as for CVB3. However, we show that this cleavage attenuates the interaction of TFEB with RagC, an interaction that promotes TFEB trafficking to the surface of the lysosomes (16). Intriguingly, despite this disruption of the TFEB-RagC interaction, TFEB continues to localize to the cytosol during EV-D68 infection. We further demonstrate that overexpression of a mutant TFEB construct (Δ30-TFEB) lacking the RagC binding domain blocks autophagic flux and increases EV-D68 egress, while overexpression of wild-type TFEB (WT-TFEB) does not. Our results suggest that TFEB plays multiple roles in EV-D68 infection by promoting viral RNA replication and autophagosome-mediated nonlytic release of infectious EV-D68 particles.

## Results

### TFEB is essential for EV-D68 genomic RNA replication

TFEB has recently been shown to be essential for Coxsackievirus egress from cells, but whether the cellular transcription factor is required for EV-D68 infection is unknown (17). To determine whether TFEB is necessary for EV-D68 infection, we knocked down TFEB (**Figure 1A**) and infected the cells with EV-D68 for 5h. As shown in **Figure 1B**, the knockdown of TFEB significantly reduces EV-D68 cell-associated titers, suggesting that the transcription factor is essential for EV-D68 infection. We wanted to understand how TFEB regulates EV-D68 infection and identify the specific step in the viral life cycle that TFEB targets. For this reason, we begin by examining the effect of TFEB depletion on the early phases of the EV-D68 lifecycle, including binding and entry/fusion. As demonstrated in **Figures 1C** and **1D**, the knockdown of TFEB does not significantly affect EV-D68 binding or entry/fusion in H1HeLa cells. We then examined the effect of TFEB knockdown on EV-D68 genomic RNA replication. Scramble and TFEB knockdown cells were infected for 5h before being subjected to qPCR analysis. As shown in **Figure 1E**, knockdown of TFEB reduces EV-D68 RNA replication. Together, these results suggest that TFEB specifically regulates EV-D68 genomic RNA replication but is dispensable for EV-D68 entry.

**Figure 1.**
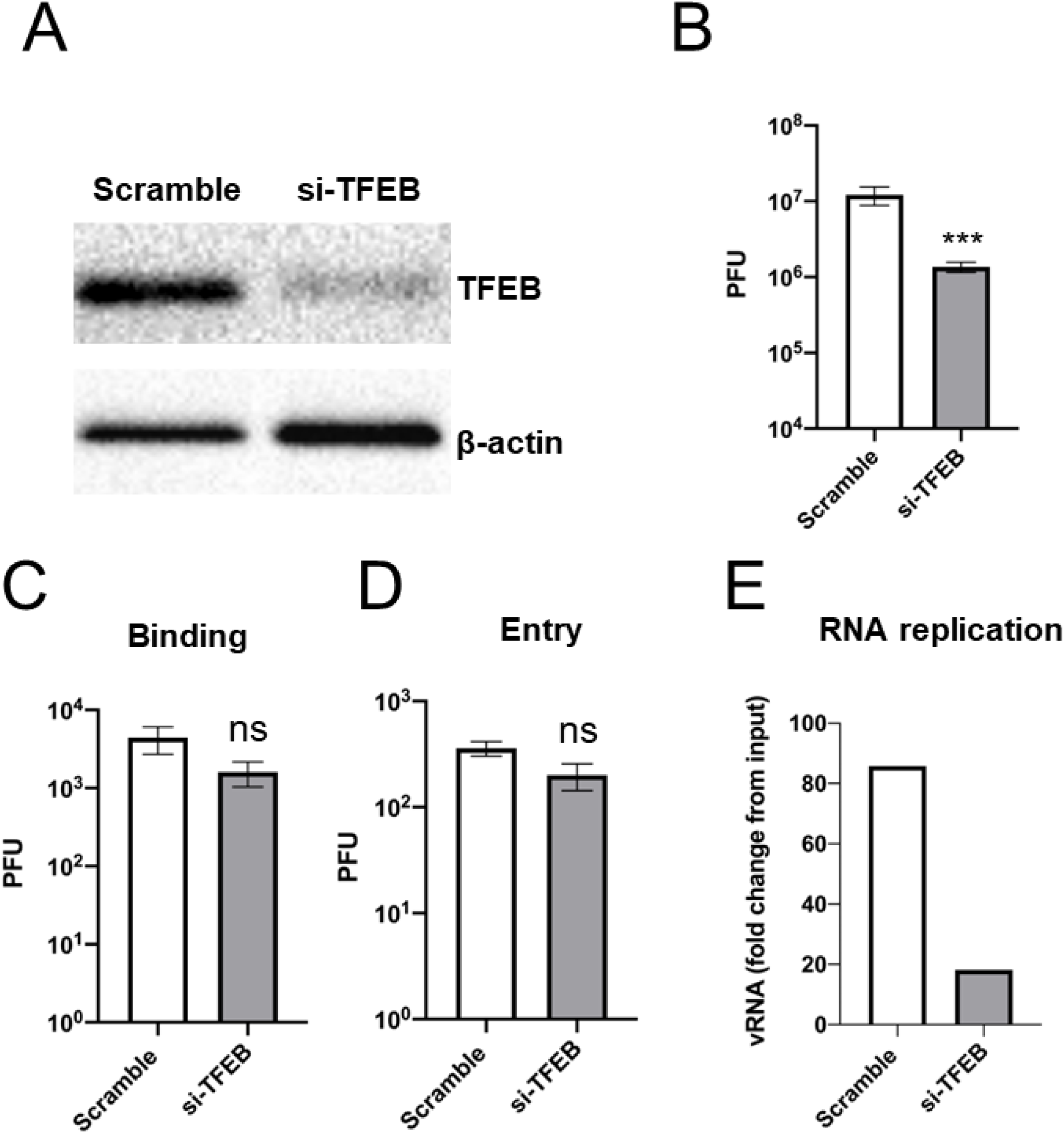
TFEB promotes EV-D68 RNA replication. (A) H1HeLa cells were transfected with the scramble control or TFEB siRNAs for 48 h. The lysates were collected and prepared for a western blot analysis. (B) Cells were transfected as in A and infected with EV-D68 (MOI =0.1) for 5 h, and viral titers were determined by a plaque assay. (C) H1HeLa cells were transfected with the scramble control or TFEB siRNAs for 48 h. The cells were prechilled on ice for 1 h and infected (MOI 30) for 1 h on ice. The cells were then collected and prepared for plaque assay. (D) Cells were transfected and prechilled as in A, then infected (MOI =30) for 30 mins on ice. The cells were washed and incubated at 37°C for 1h, then prepared for a plaque-assay-based viral titer determination. (E) H1HeLa cells were transfected as in A and infected for 5 h for RNA isolation and cDNA synthesis. Viral RNA replication was determined by qPCR. Error bars indicate mean ± SEM of at least 3 independent experiments. Unpaired student’s t-test was used for the statistical analysis (***= p< 0.001; ns=not significant.).

### TFEB does not colocalize with dsRNA, and its subcellular localization is not altered during EV-D68 infection

Since the knockdown of TFEB specifically reduced EV-D68 genomic RNA replication and to understand how TFEB promotes EV-D68 RNA replication, we examined whether TFEB’s subcellular localization changes during EV-D68 infection and whether the transcription factor colocalizes with dsRNA, a viral RNA replication intermediate, used to demarcate viral RNA replication sites. As shown in **Figure 2**, the starvation control induced extensive TFEB relocalization to the nucleus as expected, while EV-D68 infection did not significantly alter TFEB’s subcellular localization. We then examined whether TFEB colocalizes with dsRNA. Although TFEB and dsRNA appear proximal to each other, we did not observe any significant colocalization between the TFEB and dsRNA signals, indicating that TFEB is not likely to associate with the EV-D68 replicase (**Figure 2**).

**Figure 2.**
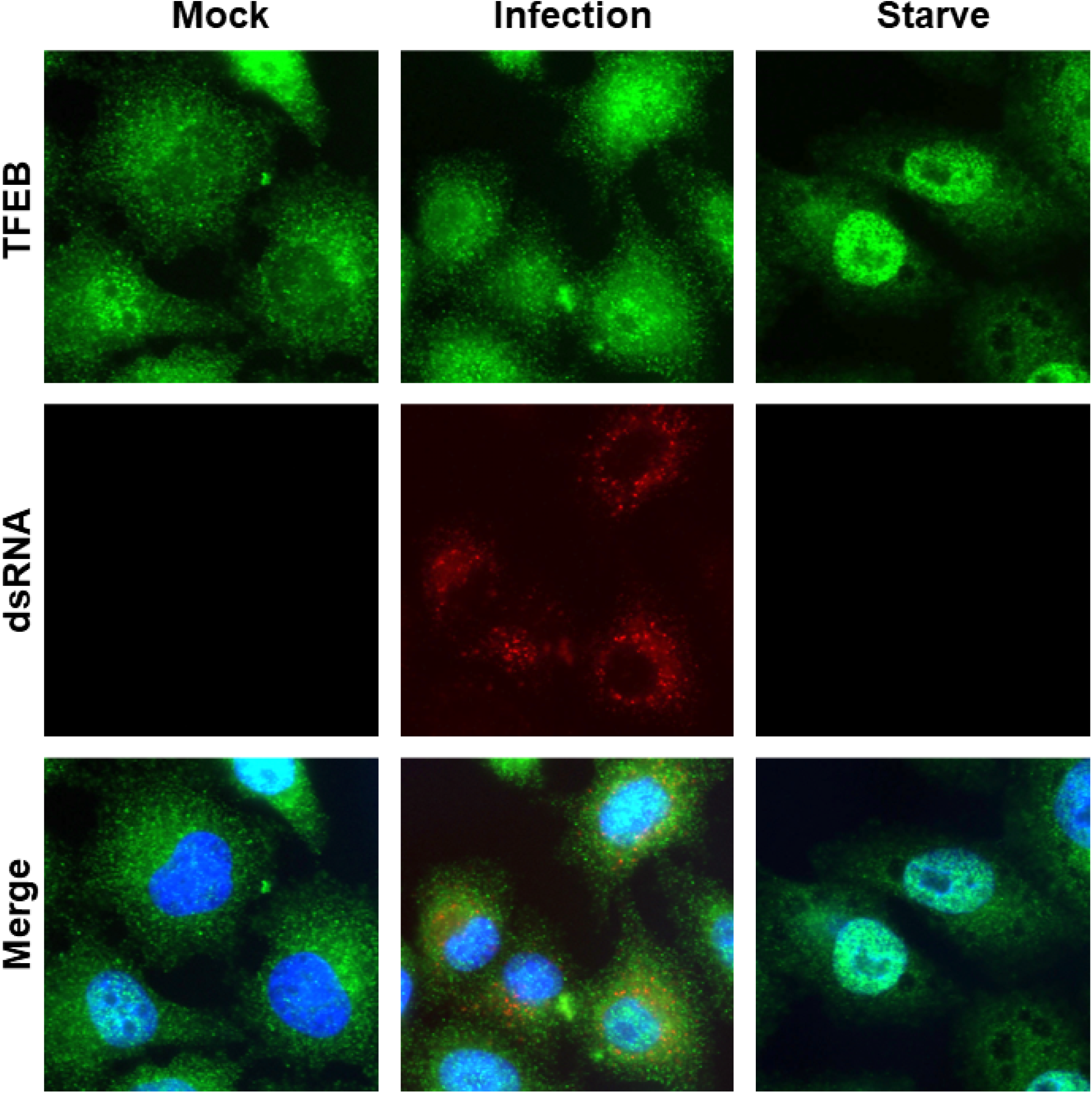
EV-D68 infection does not significantly alter TFEB’s subcellular localization. H1HeLa cells were infected with EV-D68 (MOI =30) for 4 h. The cells were fixed and immunofluorescence microscopy was performed against TFEB and dsRNA.

### The 3C protease of EV-D68 cleaves TFEB at the N-terminus

A recent study demonstrated that CVB3 cleaves TFEB during infection (17). To determine whether EV-D68 infection also causes TFEB cleavage, we infected H1HeLa cells for 4h, followed by a western blot. Consistent with the above study, we observed the full-length TFEB band disappear and the appearance of a smaller band at approximately 55 kDa (**Figure 3A**). Next, we performed a time-course infection to determine the post-infection timing of TFEB cleavage. As demonstrated in **Figure 3B**, TFEB cleavage begins as early as 3hpi. By 4hpi, the full-length TFEB band has disappeared, indicating complete cleavage of TFEB immediately after the expected peak of viral RNA replication. The 3C protease carries out CVB3 cleavage of TFEB. To confirm whether the cleaved product observed during viral infection is due to EV-D68 3C-mediated cleavage of TFEB, we incubated H1HeLa cell lysates with recombinant EV-D68 3C protease. We found the appearance of a cleavage fragment of a similar size to those observed during EV-D68 infection in lysates incubated with the recombinant 3C protease, indicating that the 3C protease of EV-D68 also cleaves TFEB (**Figure 3C**). These data suggest that the viral 3C-mediated cleavage of TFEB may be conserved in enteroviruses.

**Figure 3.**
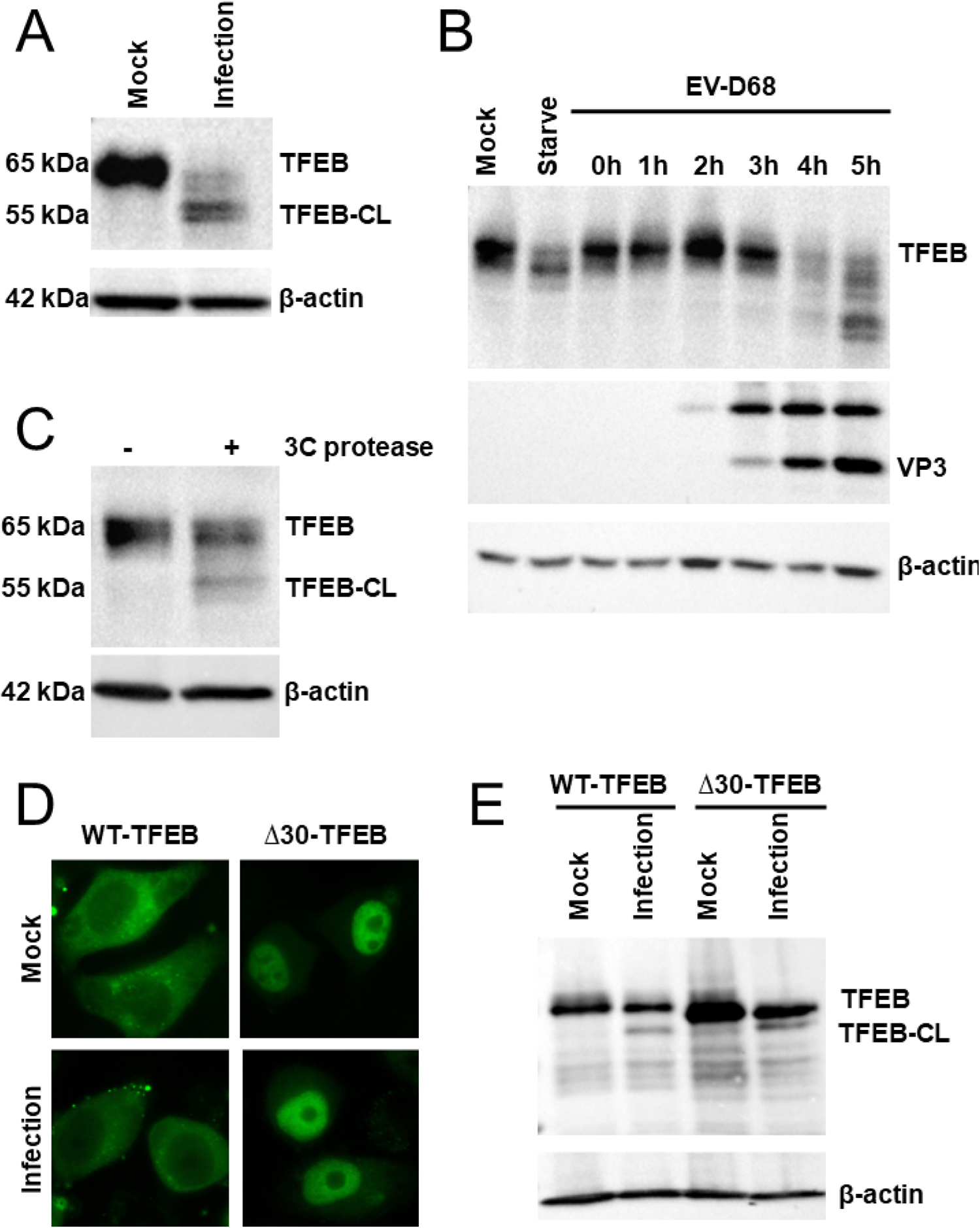
The 3C protease of EV-D68 cleaves TFEB at the N-terminus. (A) H1HeLa cells were infected with EV-D68 for 4 h. The lysates were collected at the end of the infection, and a western blot was performed against TFEB. (B) Cells were infected for the indicated time points for western blot against the indicated proteins. Starvation for 4 h was included as a control. (C) H1HeLa cell lysates were incubated with or without a recombinant 3C protein overnight at 4°C for a western blot against TFEB. (D and E) TFEB constructs (WT-TFEB and Δ30-TFEB) were expressed in H1HeLa cells for 24 h. The cells were left uninfected (mock) or infected with EV-D68 for 4 h. Images were taken using an echo revolve microscope (D) before cells were lysed for western blot (E) using anti-GFP antibodies.

To determine where the viral 3C protease cleaves TFEB, we employed our C-terminal GFP-tagged wild-type (WT-TFEB) and mutant (Δ30-TFEB) constructs. The first 30 amino acids, which correspond to the RagC binding site, are deleted in the Δ30-TFEB constructs (15). Since the TFEB-RagC interaction is essential for TFEB trafficking to the surface of the lysosomes, where it is phosphorylated and retained in the cytosol, the mutant construct is exclusively localized to the nucleus (12, 16, 18). We overexpressed these TFEB constructs in H1HeLa cells, infected them with EV-D68 (**Figure 3D**), and performed a western blot using anti-GFP antibodies (**Figure 3E**). If cleavage happens at the C-terminus, we expect to see the full-length TFEB band and a smaller fragment approximately the size of GFP. However, this is not the case, as we observed cleavage fragments of roughly 55 kDa for both constructs, suggesting that cleavage happens shortly to the C-terminal side of the RagC binding site-containing N-terminal domain (**Figure 3E**). For CVB3, the Q60/S61 site is the cleavage site, and our data are consistent with that site for EV-D68 (17).

### EV-D68 attenuates TFEB RagC interaction and TFEB trafficking to the surface of the lysosomes

Since the viral 3C protease cleaves TFEB at the N-terminus, removing the RagC binding domain, we hypothesized that TFEB-RagC interaction would be disrupted during EV-D68 infection. One expected consequence of this would be reduced trafficking of TFEB to lysosomes during infection. Consistent with this hypothesis, TFEB trafficking to the surface of the lysosomes, as measured by its colocalization with LAMP1, a lysosome marker, was reduced during EV-D68 infection relative to the mock infection control (**Figure 4A**). Despite this, however, TFEB remains largely cytosolic during EV-D68 infection (**Figure 4A**). To specifically test TFEB-RagC interaction during infection, we overexpressed WT-TFEB, immunoprecipitated for RagC, and immunoblotted using anti-GFP antibodies. We also included the Δ30-TFEB construct as a control. As shown in **Figure 4B**, EV-D68 infection diminished the amount of co-immunoprecipitated TFEB-RagC in WT-TFEB overexpressing cells. Densitometry analysis shows the ratio of TFEB/RagC in the pulldown drops from 1.04 to 0.40 during infection. As expected, our Δ30-TFEB shows weak interaction with RagC, irrespective of EV-D68 infection, with a TFEB/RagC ratio of 0.18 in mock and 0.22 in infected cells. Together these data lead us to conclude that RagC interaction with TFEB is reduced during infection.

**Figure 4.**
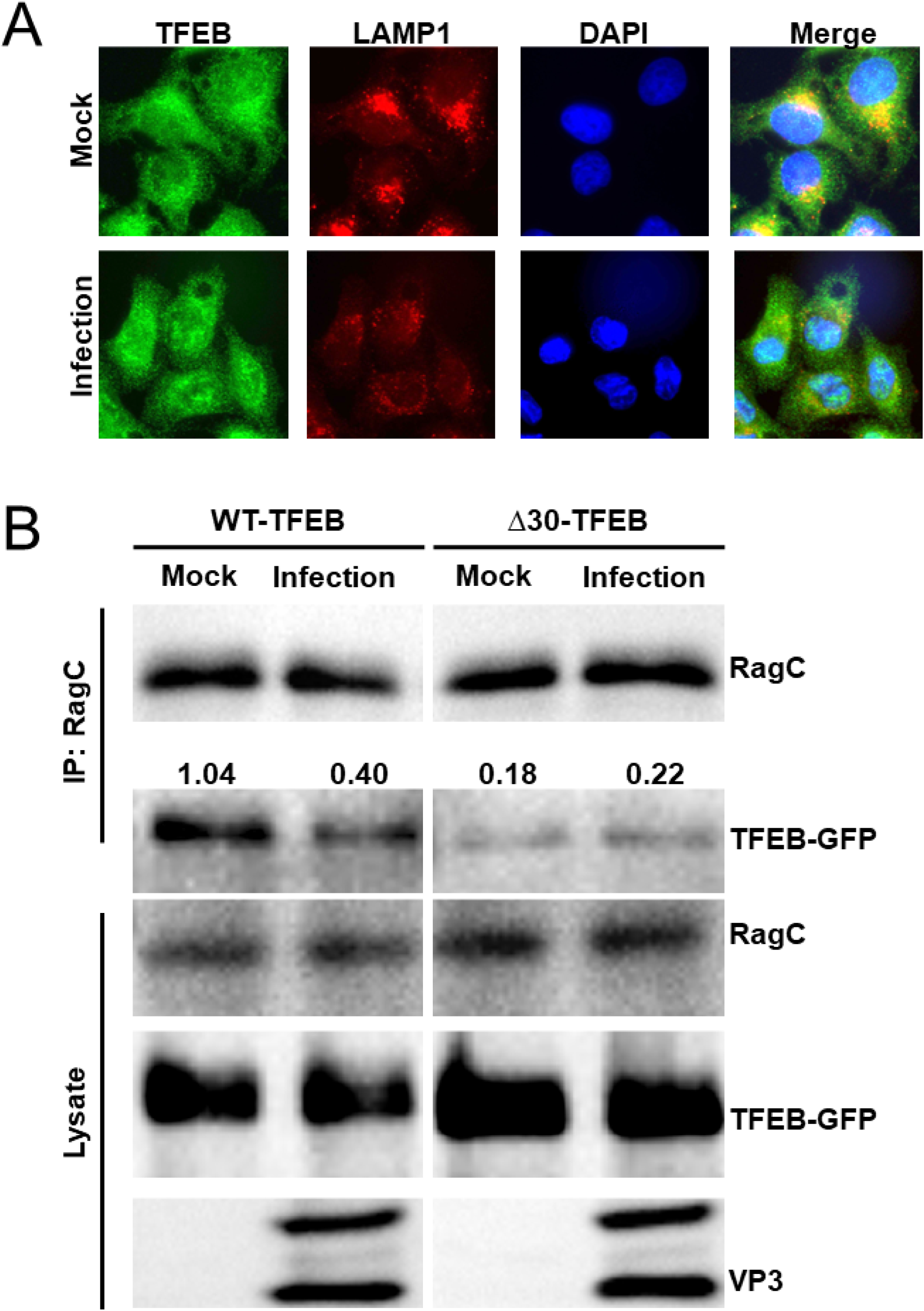
EV-D68 attenuates TFEB RagC interaction and TFEB trafficking to the surface of the lysosomes. (A)H1HeLa cells were infected with EV-D68 (MOI =30) for 4 h. The cells were fixed, and immunofluorescence microscopy was performed against TFEB and LAMP1. (B) H1HeLa cells were transfected with the WT-TFEB and Δ30-TFEB constructs for 24 h. The cells were then infected with EV-D68 (MOI =30) for 4 h, after which lysates were collected and subjected to immunoprecipitation using anti-RagC antibodies. The numbers above the TFEB blots (B) represent ratios of TFEB/RagC derived from densitometry analysis.

### Knockdown of RagC induces TFEB translocation to the nucleus and reduces EV-D68 titers

Given that RagC is required for TFEB localization to the cytoplasm, we hypothesized that knockdown of RagC will induce TFEB translocation to the nucleus, making TFEB inaccessible to the virus and consequently decreasing viral titers (16). To test this, we first knocked down RagC and performed immunofluorescence microscopy against TFEB. As expected, the knockdown of RagC causes TFEB nuclear localization while the scramble control group TFEB is mainly localized to the cytoplasm **Figure 5A**). We observed a reduction of TFEB cleavage in RagC knockdown cells compared to the scramble siRNA control during EV-D68 infection. (**Figure 5B**). Finally, we examined the effect of RagC knockdown on viral intracellular titers. RagC knockdown, similar to TFEB knockdown, significantly reduced EV-D68 intracellular titers (**Figure 5C**).

**Figure 5.**
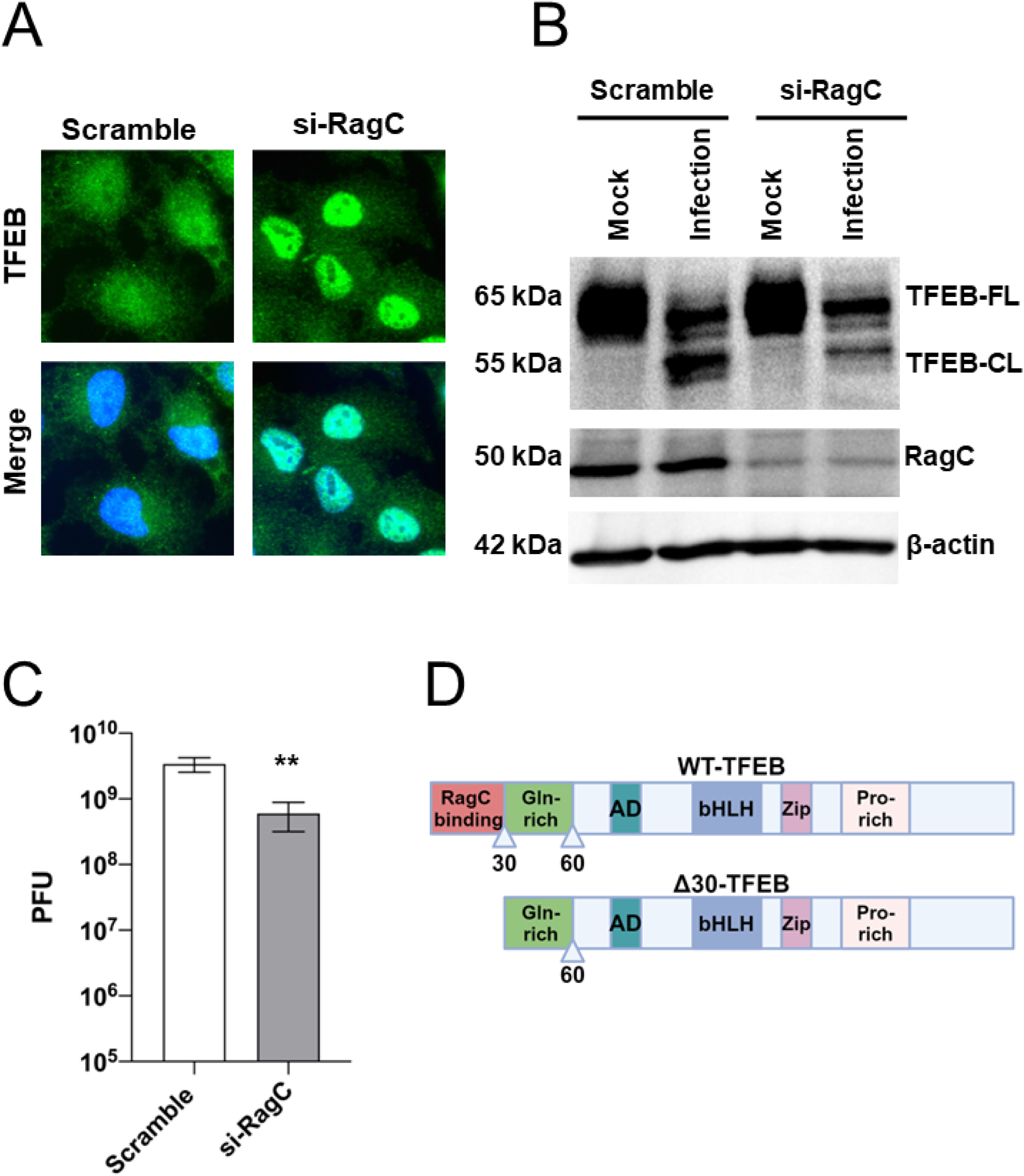
Knockdown of RagC decreases EV-D68 titers. (A) Cells were transfected with the indicated siRNAs for 48 h. The cells were then fixed, and immunofluorescence microscopy was performed against TFEB. (B) H1HeLa cells were transfected as in A. The cells were then infected with EV-D68 (MOI =30) for 4 h. Lysates were collected and prepared for a western blot against the indicated antibodies. (C) H1HeLa cells were transfected with scramble and TFEB siRNAs for 48 h, followed by 0.1 MOI EV-D68 infection for 5 h. Viral titers were measured using a plaque assay. Error bars denote the mean ± SEM of 3 independent experiments. Unpaired student’s t-test was used for the statistical analysis (**= p< 0.01). (D) Schematics of WT-TFEB and Δ30-TFEB constructs, not to scale. RagC binding domain, Glutamine-rich domain, Activation domain (AD), basic helix-loop-helix domain (bHLH), Zip domain, and Proline-rich regions are shown. Triangles represent the cutoff for Δ30-TFEB and the established cleavage site for CVB3 3C protease.

### Overexpression of Δ30-TFEB, not WT-TFEB, blocks autophagic flux and promotes EV-D68 release

Our data indicate that EV-D68 3C protease cleaves TFEB at the N-terminal region, attenuating TFEB-RagC interaction. Since the cleavage product generated during EV-D68 lacks the RagC binding domain, much like our Δ30-TFEB construct, we wanted to understand the significance of TFEB cleavage during EV-D68 infection. We overexpressed the TFEB constructs in H1HeLa cells, then infected the cells with EV-D68 for 5h. While overexpression of WT-TFEB and Δ30-TFEB constructs did not significantly impact EV-D68 intracellular titers (**Figure 6B**), overexpression of Δ30-TFEB increases EV-D68 release (**Figure 6C**) in H1HeLa cells. To tease out how Δ30-TFEB enhances EV-D68 nonlytic release, we overexpressed our TFEB constructs in cells and starved or treated them with ammonium chloride to induce or block autophagic flux. While amino acid starvation induces p62 degradation in pcDNA3.1 and WT-TFEB overexpressing cells, p62 degradation was severely attenuated in cells overexpressing Δ30-TFEB, indicating that Δ30-TFEB, which enhances EV-D68 release, also blocks autophagic flux (**Figure 6A**). These data suggest that TFEB cleavage during EV-D68 could be a mechanism the virus uses to block autophagic flux, generating a backup of autophagic vesicles for nonlytic release. If this hypothesis is correct, we reasoned that overexpressing the Δ30-TFEB construct in ATG7 KO H1HeLa cells, which cannot form autophagosomes, would not increase EV-D68 release. Consistent with this reasoning, overexpression of neither WT-TFEB nor Δ30-TFEB constructs in ATG7 KO affect EV-D68 release (**Figure 6D**).

**Figure 6.**
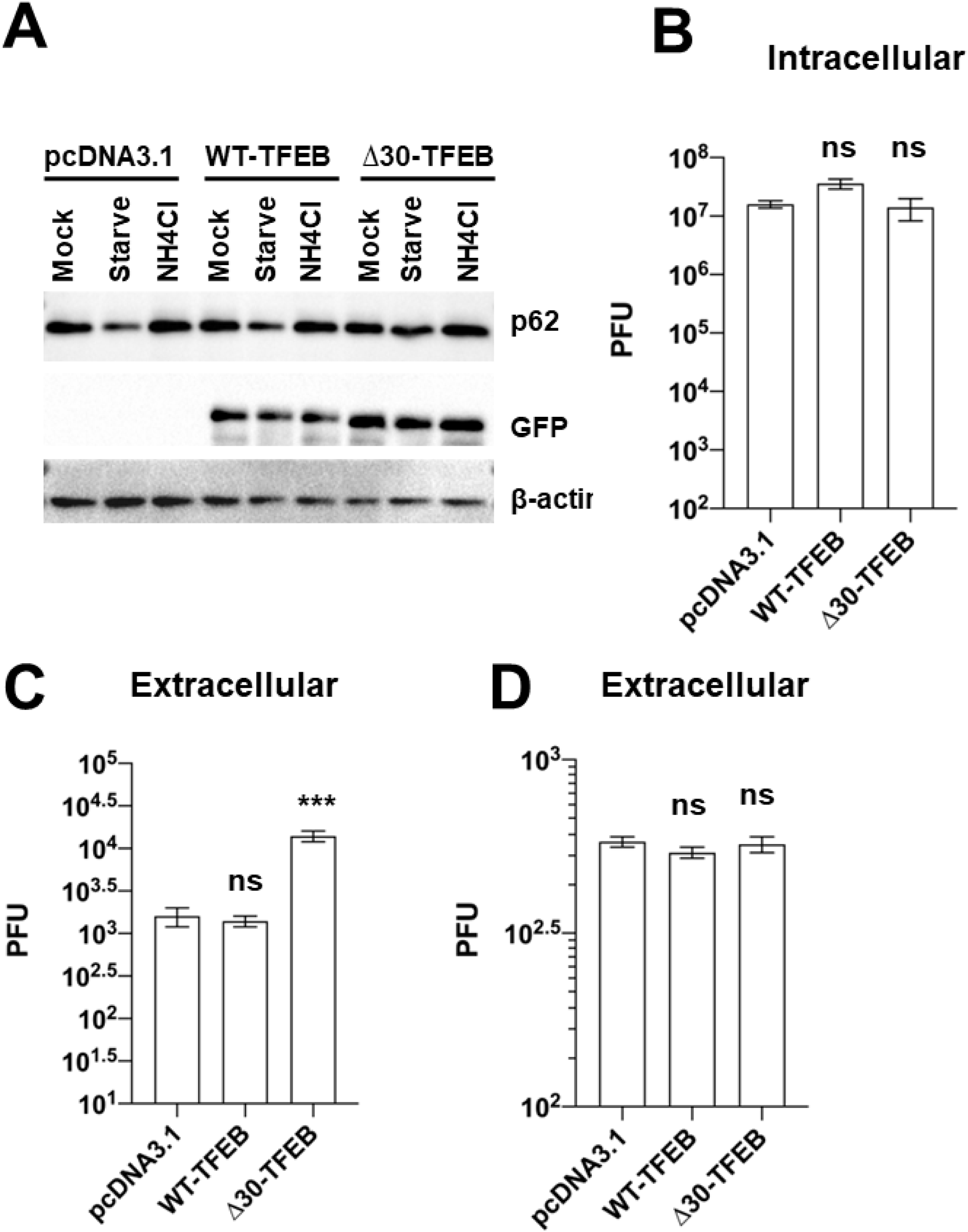
Overexpression of Δ30-TFEB, not WT-TFEB, promotes EV-D68 release. (A) Cells were transfected with the indicated plasmid constructs for 24 h. The cells were then left untreated (mock), starved for 4 h (starve), or treated with ammonium chloride for 4 h. Lysates were harvested and prepared for western blot. (B and C) cells were transfected as in A and infected with EV-D68 (MOI =0.1) for 5 h. The intracellular (B) and extracellular (C) viral titers were determined by a plaque assay. (D) ATG7 KO H1HeLa cells were transfected with the indicated plasmids for 24 h and infected as in B. Extracellular viral titers were measured by a plaque assay. Error bars indicate mean ± SEM of at least 3 independent experiments. Unpaired student’s t-test was used for the statistical analysis (***= p< 0.001; ns=not significant.).

## Discussion

Many enteroviruses are known to usurp components of the cellular autophagy pathway to resculpt membranes for their benefit. Here, we demonstrate that TFEB, the master transcriptional regulator of autophagy and lysosomal biogenesis, is required for EV-D68 infection of host cells. TFEB regulates EV-D68 in two distinct steps of the viral lifecycle using different mechanisms: genomic RNA replication and nonlytic release. Precisely how TFEB contributes to EV-D68 RNA replication is still being investigated; however, here, we show that the TFEB cleavage product generated after peak RNA replication blocks autophagic degradation and facilitates the autophagosome-mediated nonlytic release of EV-D68 particles. Our results pinpoint TFEB as a critical host factor for at least two separate steps of the EV-D68 lifecycle.

Our results show that TFEB is essential for EV-D68 RNA replication (**Figure 1B**) but not for viral entry (**Figures 1C, 1D, and 1E**). Knockdown of TFEB reduced EV-D68 RNA replication but had no major impact on viral binding or entry into host cells, suggesting that TFEB is specifically important for EV-D68 genomic RNA replication. How TFEB regulates EV-D68 genomic RNA replication is still being determined. Our immunofluorescence microscopy analysis in **Figure 2** shows little to no colocalization between TFEB and dsRNA, suggesting that TFEB does not associate with the EV-D68 replicase. Given its role in regulating autophagy, essential for enteroviral replication, we initially suspected that TFEB may promote EV-D68 RNA replication by regulating the formation of autophagosomes. However, this is not the case since TFEB knockdown in autophagy-defective ATG-7 KO cells also significantly reduces EV-D68 titers (**Figure S1**).

The 3C protease of CVB3 was recently shown to cleave TFEB during infection (17). We show here that the EV-D68 3C protease also cleaves TFEB after peak viral RNA replication. We have also observed a similar pattern of TFEB cleavage during Poliovirus infection, indicating that TFEB cleavage is a shared feature amongst many enteroviruses **(Figure S2A)**. TFEB cleavage during enteroviral infection could be significant for many reasons. First, even though EV-D68 and other enteroviruses are known to use components of autophagic signaling for their benefit, we have recently shown that stress-induced autophagy via amino acid starvation during infection (starvation after infection) impairs the infection of multiple enteroviruses, including EV-D68, CVB3, Rhinovirus-14, and Poliovirus (6). Recent unpublished data from our lab shows that TFEB regulates this stress-induced antiviral autophagy (Jassey et al. manuscript in preparation). We, therefore, hypothesized that TFEB cleavage during enteroviral infection may constitute a strategy enteroviruses use to restrict antiviral autophagy. In support of this hypothesis, we show that attenuating TFEB cleavage by knocking down RagC, which induces TFEB translocation to the nucleus, decreases viral intracellular titers (**Figure 5C** and **Figure S2B**). Secondly, TFEB cleavage appears to be an additional mechanism that enteroviruses use to block autophagic flux and ensure viral nonlytic egress. This is because overexpression of Δ30-TFEB, as shown in **Figure 6C**, but not the wild-type TFEB, blocks starvation-induced p62 degradation and increases EV-D68 nonlytic release in H1HeLa cells, but not in the autophagy defective ATG-7 KO cells (**Figure 6D**), indicating that Δ30-TFEB impairs autophagic flux and facilitates the autophagosome-mediated release of infectious EV-D68 particles, as has been previously noted for Poliovirus (19). However, precisely how Δ30-TFEB restricts autophagic flux is still being investigated.

The TFEB-RagC interaction is essential for the cytosolic retention of TFEB, where it is transcriptionally inactive (16). Our results show that the 3C protease of EV-D68 cleaves TFEB at the N-terminus, generating a cleavage product similar to our Δ30-TFEB construct, which has the RagC binding domain deleted (**Figure 3E**). In agreement with these results, we found that TFEB-RagC interaction and its colocalization with LAMP1 are severely disrupted during EV-D68 infection (**Figure 4**). To our surprise, despite attenuating TFEB-RagC interaction, which should cause TFEB to localize to the nucleus, TFEB remained localized to the cytosol during EV-D68 infection (**Figures 2, 3D,** and **4A**). This finding is different from the CVB3 paper cited above, which shows TFEB translocation to the nucleus late during infection, indicating that CVB3 and EV-D68 may use TFEB somewhat differently. The disparity between EV-D68 and CVB3 regarding TFEB localization during infection is unsurprising since we have recently shown that the deacetylase SIRT-1, which deacetylates TFEB, is essential for EV-D68 release but not for CVB3 release (20). These findings suggest that EV-D68 and CVB3 can engage the same cellular process differently. Given that inducing TFEB nuclear localization through RagC knockdown attenuates EV-D68 titers, we speculate that EV-D68 may employ a yet unknown mechanism to prevent TFEB nuclear import in the absence of RagC interaction.

Our data show that RagC and TFEB are also essential for Poliovirus infection. Knockdown of the cellular proteins reduced PV intracellular titers, similar to their effect on EV-D68 infection. These findings suggest that TFEB and RagC could be essential for many enteroviruses, making them attractive targets for developing broad-spectrum anti-enteroviral therapies, even if the mechanism of action may be distinct from one member of the virus family to the other.

## Materials and Methods

### Cell culture and plasmids

H1HeLa cells were cultured in DMEM supplemented with 10% heat-inactivated fetal bovine saline, 1x penicillin-streptomycin, and 1x sodium pyruvate (Gibco, 11360-070) and incubated in a 5% CO_2_ incubator at 37°C. The TFEB wild-type (WT-TFEB [plasmid #38119]) and mutant (Δ30-TFEB [plasmid #44445]) constructs were obtained from Addgene (deposited by Shawn Ferguson) and transfected into H1HeLa using the Lipofectamine 2000 transfection reagent. The transfection complex was replaced with basal growth media 6 h post-transfection, and viral infections were initiated 24 h post-transfection.

### Western blotting

H1HeLa cells plated in 6-well plates were infected and lysed at the end of the infection with RIPA buffer supplemented with cOmplete Tablets Mini Protease Inhibitor Cocktail. Lysates were collected and kept on ice for at least 30 minutes and clarified at 14000 rpm for 30 minutes. The clarified lysates were transferred into Eppendorf tubes, and protein concentrations were determined by a Bradford assay. Equal amounts of lysates were run on SDS-PAGE before being transferred to polyvinylidene fluoride membranes. Blocking was performed at room temperature on a shaker for 1 h with 5% skim milk, after which the membranes were washed twice (10 minutes per/wash) with TBST (100 mM Tris hydrochloride, pH 7.4, 2.5 M sodium chloride, and 0.125% Tween-20), and incubated with primary antibodies overnight on a shaker at 4°C. The membranes were again washed as described above and incubated with HRP-conjugated secondary antibodies for 1 h at room temperature. Western Lightning ECL was then used to develop the membranes before image acquisition using ChemiDoc (Bio-Rad).

### Viral Infections

Viral infections were performed with either low MOI (MOI of 0.1 for 5 h) or high MOI (MOI 30 for 4 h), as previously described. Briefly, viral particles were diluted in serum-free media, added to cells, and incubated at 37°C for 30 minutes. The cells were washed twice with PBS, overlaid with growth media, and incubated at 37°C until the end of the infection.

### Plaque assays

For intracellular viral titers, viral particles collected after three freeze-thaw cycles were serially diluted in serum-free media, added to cell monolayers, and incubated for 30 minutes at 37°C. For extracellular viral titer measurement, supernatants collected after viral infection were serially diluted above. The cells were then overlaid with a 1:1 ratio of 2x MEM and 2% agar, and plaques were developed 48 h after the agar overlay.

### siRNA Knockdowns

For siRNA-mediated knockdown of TFEB and RagC, H1HeLa seeded in 6-well plates to 40% confluency were transfected with 200 nM of the target siRNAs and the scramble control siRNA using Opti-MEM transfection media and Lipofectamine 2000. The transfection complex was removed 6 h post-transfection and replaced with growth media. Knockdown efficiency and viral infections were performed 48 h post-transfection.

### RNA isolation and quantitative polymerase chain reaction (qPCR)

TRIzol was used to isolate total RNA from cells according to the manufacturer’s instructions. Thermo Scientific RevertAid H Minus First Strand cDNA Synthesis Kit (K1632) was used to synthesize the cDNA after genomic DNA removal. qPCR was conducted using KiCqStart SYBR qPCR Ready Mix (Sigma, KCQS01) with the 7500 Fast Dx Real-Time PCR Instrument (Applied Biosystems) according to the manufacturer’s protocols. The following primers, 5′ TAACCCGTGTGTAGCTTGG-3′ and 5′ -ATTAGCCGCATTCAGGGGC-3′, which are specific to the 5′ untranslated region (UTR), were used to amplify EV-D68, and gene expression results were normalized to GAPDH and plotted as relative expression compared to the input virus.

### Immunofluorescence analysis

Cells were fixed with 4% PFA for 20 minutes and permeabilized with 0.3% Triton-X for 30 minutes. The cells were blocked with 3% bovine serum albumin for 1 h on a shaker. The primary antibodies diluted in the blocking buffer were added to the cells and incubated overnight on a shaker at 4°C. The cells were washed thrice with PBS, and the secondary antibodies were added and incubated at room temperature for 1 h. The cells were again washed three times before being stained with a nuclear stain (Dapi) and imaged using a revolve microscope.

### Immunoprecipitation

Immunoprecipitation was performed using protein A magnetic beads from cell signaling. Cells were lyzed using NP-40 lysis buffer and incubated on ice for at least 30 minutes. The clarified lysates were precleared and incubated overnight with anti-RagC antibodies in the cold room. The protein A magnetic beads were then added to the lysate-antibody complex and incubated at room temperature with rotation for 30 minutes. The beads were washed three times with the NP-40 lysis buffer, eluted with the 5X loading dye, and cooked at 95°C for 10 minutes.

### Virus entry assays

Viral entry assays were conducted as previously described. For the viral binding assay, H1HeLa cells seeded in 6-well plates were prechilled on ice for 30 minutes. The cells were then infected with EV-D68 and kept on ice for 1 h. They were then collected and processed for plaque assay as described above. For viral entry, cells were prechilled and infected with EV-D68 for 30 minutes on ice to allow viral binding. They were then washed twice with ice-cold PBS and shifted to 37°C for 1 h to permit viral entry. Finally, they were collected and prepared for a plaque assay-based viral titer determination.

### Statistical analysis

GraphPad Prism software (Version 7.03) was used for statistical analyses, with values representing mean ± standard error of the mean (SEM) of at least 3 independent repeats unless otherwise indicated. Student t-test was used for comparison, and statistical significance was set at a p-value of < 0.05. Densitometry analysis was performed using the FIJI package (21).

---

**Supplemental Figure 1. Knockdown of TFEB reduces EV-D68 intracellular titers in ATG7 KO cells.** Cells were transfected with the indicated siRNAs for 48 h and then infected with EV-D68 (MOI =0.1) for 5h. After three freeze-thaw cycles, intracellular viral titers were determined using a plaque assay. n = 2 independent experiments.

**Supplemental Figure 2. TFEB and RagC are essential for PV infection.** (**A**) H1HeLa cells were infected with PV (MOI =30) for the indicated time points for western blot against the indicated antibodies. (**B**) Cells were transfected with the indicated siRNAs for 48 h. The cells were then infected with EV-D68 (MOI =0.1) for 5 h. Viral titers were determined by a plaque assay.

